# A novel qPCR-based technique for identifying avian sex: an illustration within embryonic craniofacial bone

**DOI:** 10.1101/2023.03.02.530503

**Authors:** Claire J. Houchen, Maria Bergman-Gonzalez, Erin E. Bumann

**Affiliations:** Department of Oral and Craniofacial Sciences, School of Dentistry, University of Missouri-Kansas City, US

**Keywords:** sex differences, sex typing, *HINTW* gene, RT-qPCR, duck, quail, chicken

## Abstract

Sex is a biological variable important to consider in all biomedical experiments. However, analyzing sex differences in avian models can be challenging as the sexes are morphologically indistinguishable in most avian embryos. Unlike humans, female birds are the heterogametic sex with a Z and W chromosome. The female-specific W chromosome has previously been identified using species-specific polymerase chain reaction (PCR) techniques. We developed a novel quantitative real-time PCR (RT-qPCR) technique which amplifies the W chromosome gene histidine triad nucleotide binding protein W (*HINTW*) in chick, quail, and duck. We confirmed the accuracy of the single set of *HINTW* RT-qPCR primers in all three species using species-specific PCR. Bone development-related gene expression was then analyzed by sex in embryonic lower jaws of duck and quail, as duck beak size is known to be sexually dimorphic while quail beak size is not. Trends towards sexual dimorphism were found in duck gene expression but not in quail, as expected. Our novel *HINTW* RT-qPCR technique to identify the sex of avian embryos is a useful tool for including sex as a biological variable in analysis of a variety of tissues and cells used in developmental biology research.

## Main Text

Sex is known to influence the development of human organs and organ systems, such as the skeletal system (Broere-Brown et al., 2016; Hasselstrøm et al., 2006). It is therefore critical to analyze sex as a biological variable in both clinical and preclinical developmental studies. Further, analyzing sex as a biological variable is mandated by some funding bodies, such as the National Institutes of Health (NIH), in an effort to increase reproducibility and counteract overrepresentation of male animals, tissues, and cells in preclinical research (Clayton & Collins, 2014). Birds such as the chicken (*Gallus gallus*) have long been important developmental biology model organisms, because they are experimentally convenient for observing and manipulating *in ovo* embryogenesis. A complication of using an avian developmental model is that sex is morphologically indistinguishable in most avian embryos, necessitating the use of a molecular technique to identify the sex of an avian embryo.

Humans have X and Y sex chromosomes with males being the heterogametic sex (Figure 1A), while birds have Z and W sex chromosomes with females being the heterogametic sex (Figure 1B). There are instances when presence or absence of a sex chromosome may not reflect the sex phenotype of an individual of any species, but presence of a W chromosome in birds should usually indicate a female phenotype. Determining bird sex has historically been complicated by our lack of understanding of the genetic mechanisms underlying avian sex determination, but recent evidence points to a dose-dependent Z chromosome mode of sex determination in birds (Ioannidis et al., 2021; Smith et al., 2009).

**Figure 1.**
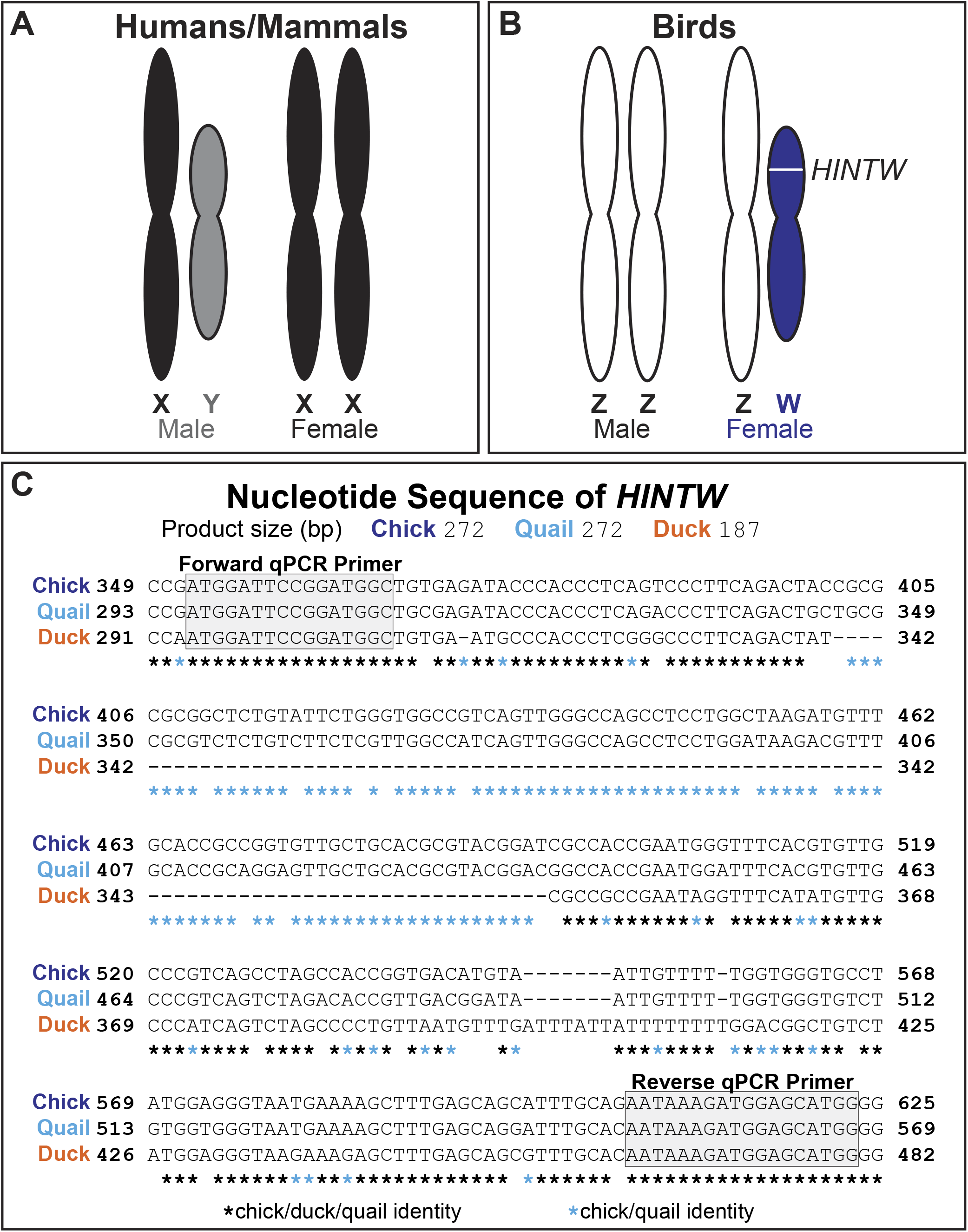
The female specific *HINTW* gene is conserved in chick, quail, and duck, and can be identified with a single novel *HINTW* RT-qPCR primer set. A) Humans have X and Y sex chromosomes with males being the heterogametic sex, while B) birds have Z and W sex chromosomes with females being the heterogametic sex. The *HINTW* gene is located on the W chromosome and thus should only be present in female birds. C) Our novel *HINTW* RT-qPCR primers align to regions of the *HINTW* gene identical in chick, quail, and duck. The RT-qPCR amplicon of these primers is 272bp in female chick and quail but is 187bp in female duck.

Histidine triad nucleotide binding protein W (*HINTW,* also sometimes called *WPKCI*) is a gene on the W chromosome (Figure 1B). *HINTW* likely does not play a role in sex determination, which aligns with the recent evidence suggesting avian sex is Z chromosome dose-dependent (Ayers et al., 2013; Ioannidis et al., 2021; Smith et al., 2009). Though likely not involved in sex determination, *HINTW* is a well-established marker of female sex in chick cells and tissues as it is absent in male cells and tissues (Clinton, 1994; Nagai et al., 2014).

It is ideal for there to be a multitude of tools to identify avian sex because commonly-used sex-typing methods tend to only work in a subset of bird species; additionally, it would be prudent in some cases to use multiple sex-typing methods to verify the sex of an organism (Dawson et al., 2016; Griffiths et al., 1998). To this end, this manuscript describes a novel quantitative real-time polymerase chain reaction (RT-qPCR) technique to amplify *HINTW* not only in chick, but also other bird models used in embryological studies such as quail (*Coturnix japonica*) and duck (*Anas platyrhynchos*). This single set of *HINTW* RT-qPCR primers identifies sex of chick, quail, and duck. Chick and quail are close evolutionary relatives and consequently have high *HINTW* sequence identity to each other and the same *HINTW* RT-qPCR product size of 272bp (Figure 1C). The *HINTW* RT-qPCR primer set produces a 187bp product for the less closely related duck (Figure 1C). We designed the forward and reverse *HINTW* primers in areas where sequences were identical in all three avian species (Figure 1C). Chick and duck sex was validated by analyzing genomic DNA using established species-specific PCR primers and quail sex was validated using newly designed quail specific *HINTW* PCR primers. All PCR and RT-qPCR primer sequences, amplicon sizes, species, and sources are listed in Table 1.

**Table 1.**
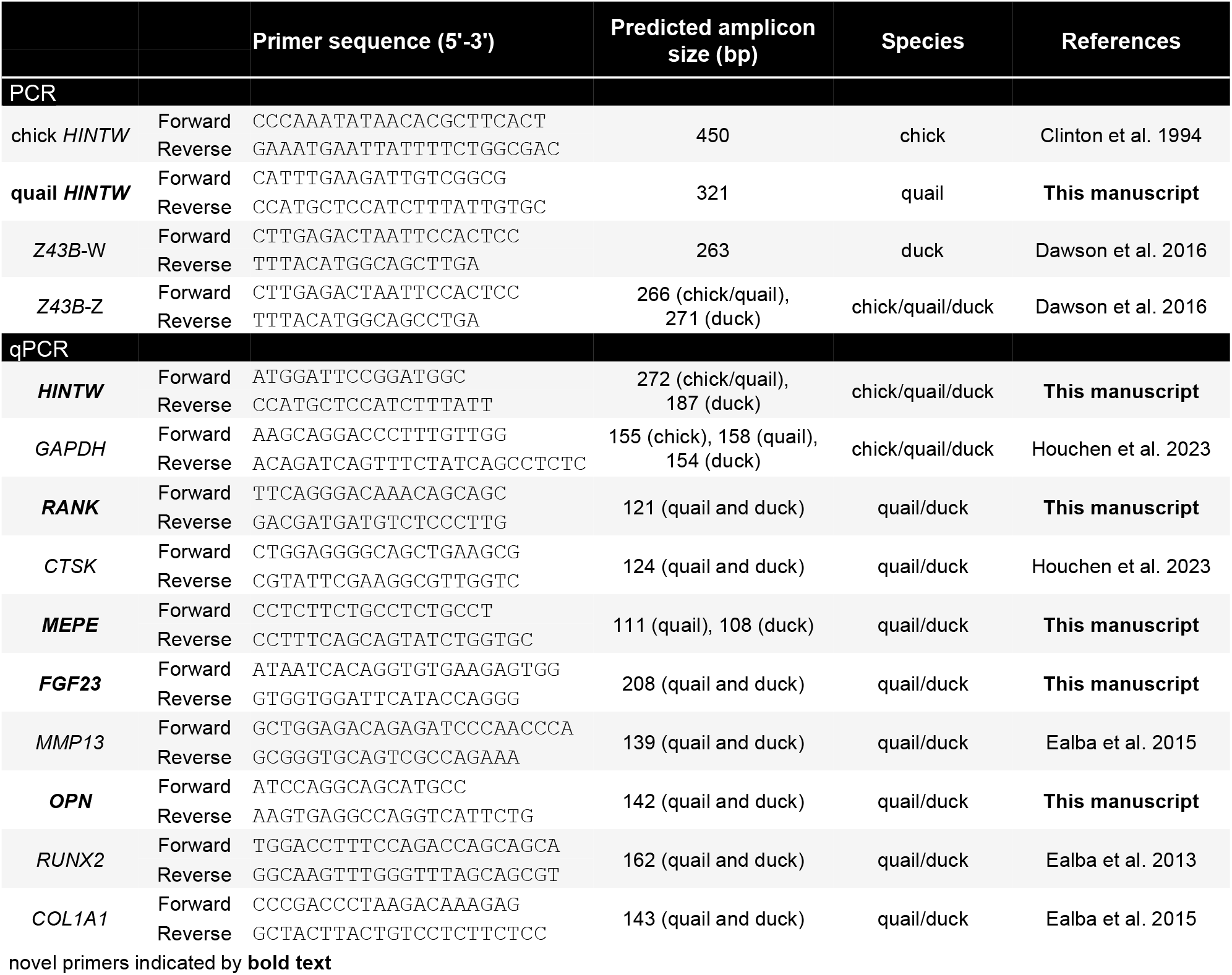
Newly designed and previously published PCR and RT-qPCR primers. Gene target, forward and reverse primer sequences, predicted amplicon size, species, and references, as appropriate, are listed. Novel primers are indicated by bold text.

Male and female chick were identified via PCR using the already-validated chick PCR *HINTW* primers (Figure 2A) (Clinton, 1994). *Z43B* primers specific to the Z chromosome (“*Z43B*-Z”) was used as an internal and technical positive control for PCR since it is present in both males and females (Figure 2A). Messenger RNA from those same individuals was tested via RT-qPCR using our novel *HINTW* primer. Female chick *HINTW* amplified at approximately 20 cycles, but products in male chick were undetected or amplified after cycle 35, which was considered a negative result (Figure 2B). The chick *HINTW* RT-qPCR product in females was 272 bp long and melted at approximately 89°C; no male RT-qPCR product was detected by melt curve analysis (Figure 2C) or gel electrophoresis (Figure 2D). For RT-qPCR, melt curve analysis, and gel electrophoresis, *GAPDH* served as an internal and technical positive control. This suggests our novel *HINTW* RT-qPCR primer reliably sex-types chick embryos.

**Figure 2.**
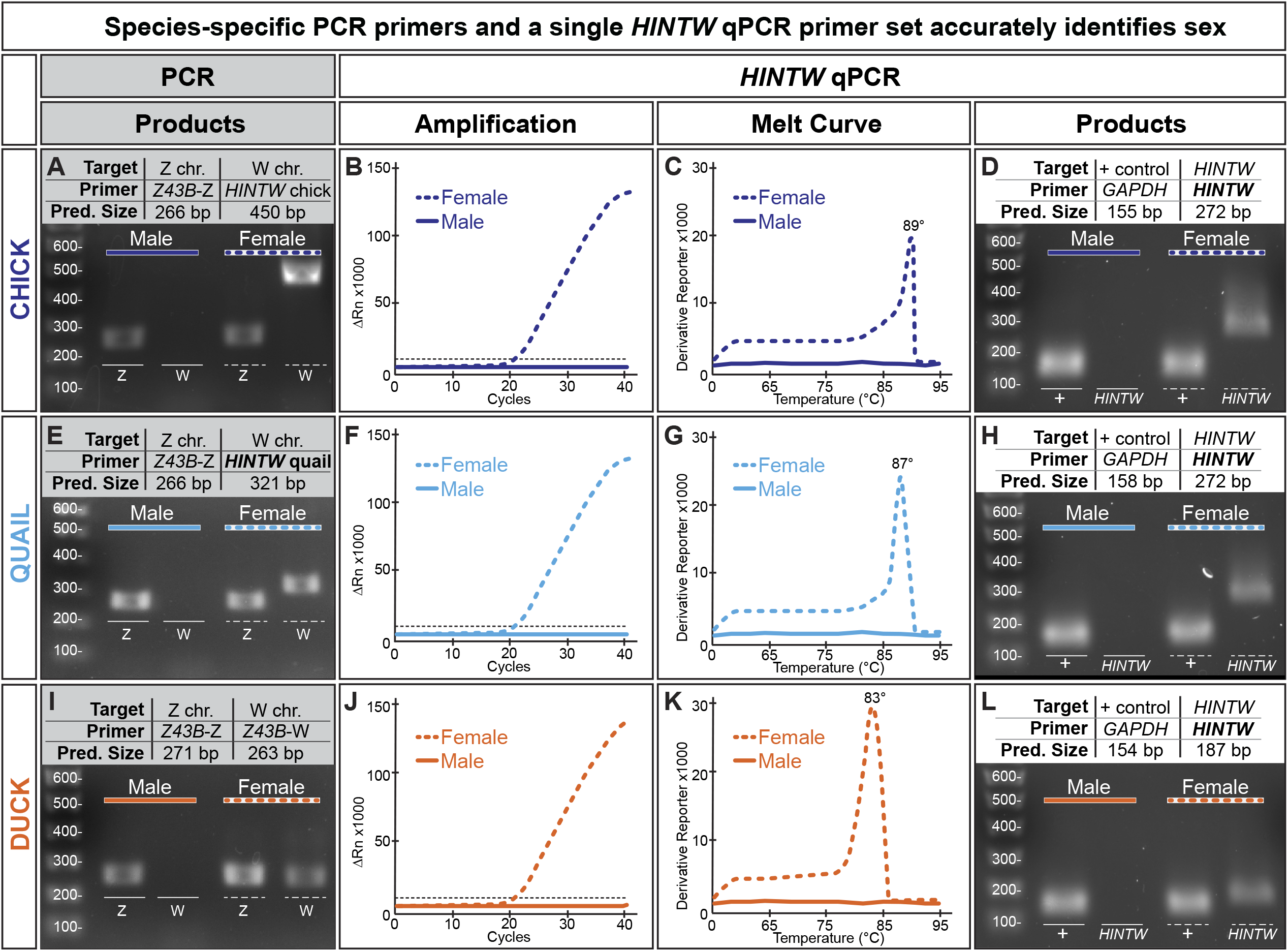
Species-specific PCR primer sets and a single *HINTW* RT-qPCR primer set accurately identifies sex. A) Chick (in dark blue) DNA was analyzed by PCR using the Z chromosome-specific *Z43B*-Z primer set and the W chromosome-specific *HINTW* chick-specific primer set. Homogametic males display a Z chromosome band but no W chromosome band, while heterogametic females display both a Z and W chromosome band. B) Messenger RNA was extracted from the same chick and was analyzed by RT-qPCR using our novel *HINTW* RT-qPCR primer set. Female chick RNA amplified around cycle 20, while male chick RNA did not amplify or amplified after cycle 35, which is considered a negative result. C) The melt curve summary graph displays a single 89°C melt peak of the *HINTW* product in females. D) RT-qPCR products were run on a gel to compare predicted amplicon size to actual size. As indicated by amplification and melt curve analysis, no product was present in male chick while there was a single 272bp female chick amplicon. *GAPDH* was the positive control. E) Quail (in light blue) DNA was analyzed using the *Z43B*-Z primers and our novel quail-specific *HINTW* PCR primers. As in chick, both sexes had a Z chromosome band, but only females had a W chromosome band. F) mRNA from the same quail was analyzed with our novel HINTW RT-qPCR primers. Female quail amplified around cycle 20, while males had a negative result. G) The melt curve summary graph demonstrates *HINTW*’s single 87°C melt peak in female quail, and H) the gel of the RT-qPCR products shows the single *HINTW* amplicon was present in females only. I) Similarly, duck (in orange) DNA was analyzed using the *Z43B*-Z and W chromosome-specific *Z43B*-W primers. Males and females had Z chromosome bands, but only females had a W chromosome band. J) Duck mRNA was analyzed by RT-qPCR using the *HINTW* qPCR primer set. Female duck amplified around cycle 21, while male duck had a negative result. K) The melt curve summary graph demonstrates the single 83°C melt peak of the *HINTW* product in females. L) As in chick and quail, gel electrophoresis of RT-qPCR products shows only female duck had a single *HINTW* amplicon.

To the best of our knowledge, no quail *HINTW* PCR primers have been published. Neither the chick *HINTW* nor the duck *Z43B*-W PCR primers successfully identified sex in quail samples (data not shown). In the absence of an optimal established PCR-based method for sex-typing embryonic quail tissue, we designed a novel quail *HINTW* PCR primer set which amplified a 321bp segment of the *HINTW* gene in females but did not produce a product in males (Figure 2E). RNA from male and female individuals was analyzed via RT-qPCR using our novel *HINTW* primer set. Female quail amplified approximately at cycle 20, while male quail amplified after cycle 35 or did not amplify; this was consistent with what we predicted based on the chick data (Figure 2F). The quail RT-qPCR product in females was 272bp long with a melt peak at 87°C, and no qPCR product was detected male samples (Figure 2G & H). The same positive control PCR and RT-qPCR primers were used for quail that were used for the chick PCR and RT-qPCR experiments. In addition to their chick evolutionary neighbor, these data suggest our novel *HINTW* RT-qPCR primer reliably sex-types quail embryos.

Male and female duck were identified via PCR using already-validated primers identifying the *Z43B* marker on the W chromosome (*“Z43B-W”*) (Figure 2I) (Dawson et al., 2016). Messenger RNA of duck embryos was extracted and converted to cDNA and analyzed by RT-qPCR using our *HINTW* primer set. Female duck *HINTW* amplified at approximately cycle 21, whereas male duck amplified after cycle 35 or did not amplify (Figure 2J). The *HINTW* RT-qPCR product in female samples was 187bp long and melted at 83°C, while no RT-qPCR product was detected in male samples (Figure 2K & 2L). The PCR and RT-qPCR positive control primers used for duck were the same as were used for quail and chick. These data indicate that our novel RT-qPCR *HINTW* primer successfully sex-types chick, quail, and duck.

We applied our *HINTW* RT-qPCR sexing technique to identify the sex of embryonic quail and duck using mRNA from the lower jaw. Duck beak size is known to be sexually dimorphic, while quail beak size is not (Nudds & Kaminski, 1984). Beak shape and size is constructed from the underlying bone, therefore sexual dimorphism in duck beak length could be reflected in differential expression of genes related to jaw bone development in males and females.

Having established sex of all quail and duck individuals using lower jaw mRNA, expression of genes relevant to jaw development were then analyzed and compared across sexes. Multiple cell types contribute to bone formation and lower jaw bone length in quail and duck, including osteoclasts, osteocytes, and osteoblasts (Ealba et al., 2015; Houchen et al., 2023). Osteoclasts resorb bone and express genes such as receptor activator of nuclear factor κ B (*RANK*) and cathepsin K (*CTSK*). Osteoblasts, on the other hand, build bone and differentiate into osteocytes, which are cells that are housed in mineralized bone and use mechanical sensing to regulate the homeostasis of bone. Genes enriched in osteocytes include matrix extracellular phosphoglycoprotein (*MEPE*) and fibroblast growth factor-23 (*FGF23*) (Agoro et al., 2023; Dallas & Bonewald, 2010). Other osteoblast lineage cells play a role in bone development and function; pre-osteoblasts express genes such as matrix metalloproteinase 13 (*MMP13*) and osteopontin (*OPN*). Additionally, genes such as runt-related transcription factor 2 (*RUNX2*) and collagen type I alpha 1 chain (*COL1A1*) are expressed in osteolineage cells both at the osteoblast stage and differentiated osteocyte stage (Agoro et al., 2023). These eight genes constitute only a fraction of the genes involved in bone development, and were selected for this analysis as they represent multiple cell types known to be essential to bone development, as well as represent various stages of differentiation.

We analyzed expression of genes related to bone development in lower jaw tissue at Hamburger-Hamilton (HH) stage 39, when the jaw is largely calcified and there is active bone resorption by osteoclasts, relative to expression at stage HH33, when bone deposition by osteoblasts is just beginning and there are no active osteoclasts in the lower jaw bone (Ealba et al., 2015; Hamburger, 1951; Houchen et al., 2023). Osteoclast *RANK* expression was mildly but significantly higher in male duck compared to female duck (n=3-4/stage/sex, p=0.03; Figure 3A). While osteoclast *CTSK* expression was not dimorphic (Figure 3B), osteocyte *MEPE* expression was also significantly higher in male duck than female duck (p=0.03; Figure 3C). Expression of another osteocyte gene, *FGF23*, was not statistically significantly different in male and female duck but might become so with increased statistical power (Figure 3D). Genes expressed in pre-osteoblasts and osteolineage cells, such as *MMP13*, *OPN*, *RUNX2*, and *COL1A1*, did not differ between male and female duck (Figure 3E-3H). These data suggest there might be sexual dimorphism in duck lower jaw development underlying the beak sexual dimorphism seen in adult duck.

**Figure 3.**
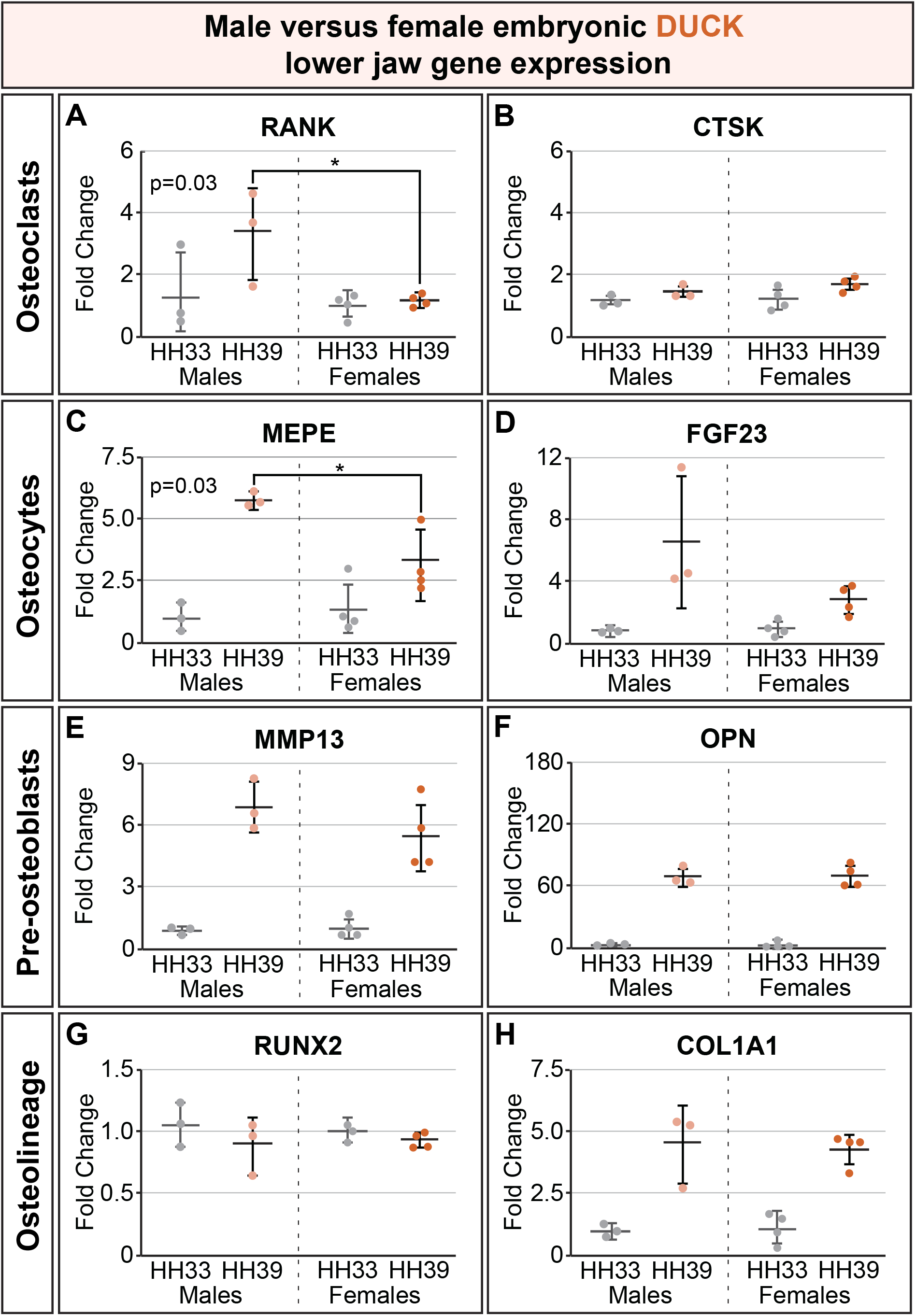
Expression of genes involved in embryonic bone development indicates possible sexual dimorphism in duck. A & B) Expression of *RANK*, a gene enriched in osteoclasts, differed significantly between male and female duck (p=0.03), but another gene expressed in osteoclasts *CTSK* did not. C) *MEPE* expression is enriched in osteocytes; *MEPE* expression also differed significantly between male and female duck (p=0.03). D) Another osteocyte gene, *FGF23* expression was not statistically significantly different in males and females but might become so with increased statistical power. E) *MMP13* and F) *OPN* are genes with enriched expression in pre-osteoblasts, and neither indicate sexually dimorphic gene expression. Similarly, G) *RUNX2* and H) *COL1A1* are genes expressed in osteolineage cells both at the osteoblast stage and at the osteocyte stage, and neither indicate differential expression in males and females. The mild sexual dimorphism indicated by these data aligns with known sexual dimorphism in adult duck beaks. *p<0.05, n=7/stage/species

Gene expression related to bone development in the lower jaw was similarly analyzed in embryonic quail at HH33 and HH39. None of the genes analyzed (*RANK*, *CTSK*, *MEPE*, *FGF23*, *MMP13*, *OPN*, *RUNX2*, and *COL1A1*) were sexually dimorphic in quail (Figure 4A-4H). This aligns with there being no documented sexual dimorphism in adult quail beaks. Gene expression analysis in Figures 3 and 4 included n=3-4 individuals/sex/species/stage and is meant to exemplify the utility of the single set of *HINTW* RT-qPCR primers to sex quail and duck embryos. Increasing N for the gene expression analyses would increase the statistical power of these tests, and our data depict trends towards presence or absence of sexual dimorphism in quail and duck jaw development gene expression rather than conclusive proof of sex differences.

**Figure 4.**
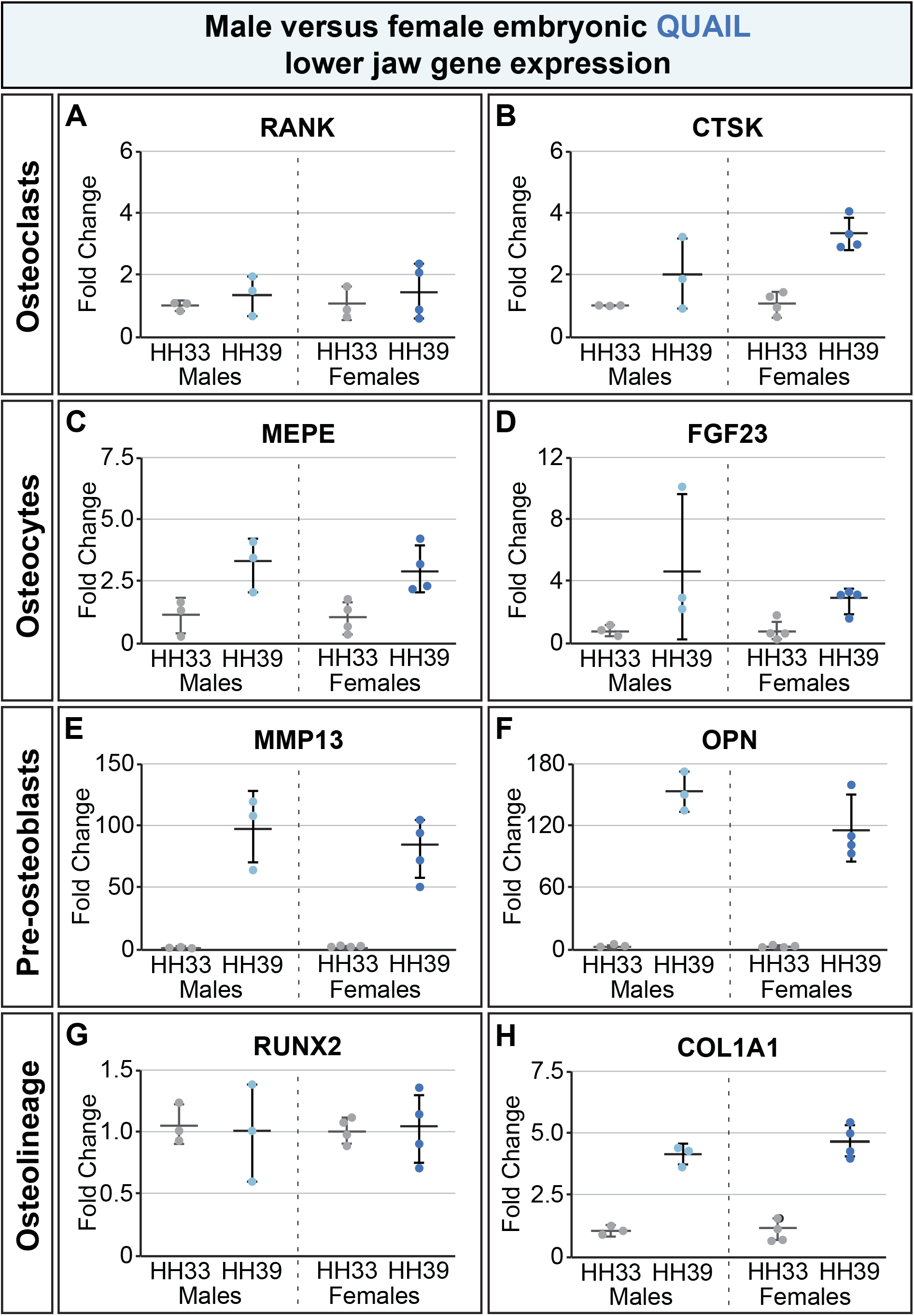
Expression of genes involved in embryonic bone development does not signify sexual dimorphism in quail. Though they represent multiple bone-relevant cell types, none of the genes analyzed (*RANK*, *CTSK*, *MEPE*, *FGF23*, *MMP13*, *OPN*, *RUNX2*, and *COL1A1*) were sexually dimorphic in quail (A-H). This reflects that in contrast to duck, quail skeletons and beaks are also not sexually dimorphic in adult specimens. *p<0.05, n=7/stage/species

In summary, our novel RT-qPCR *HINTW* technique can effectively identify sex of avian embryos in several species using a single primer set, which provides an RNA-based strategy for accomplishing the sometimes-challenging task of sex-typing birds. Additionally, this RNA-based sexing technique facilitates consideration of sex as a variable both before conducting expensive RNA analysis (such as RNA-seq) and during data analysis in studies using birds. We demonstrated the utility of this tool for considering sex as a biological variable by analyzing expression of genes related to bone development in embryonic quail and duck and found there might be sexual dimorphism in duck lower jaw development underlying the sexual dimorphism seen in adult duck beak length.

## Methods

### Use of avian embryos

Fertilized eggs of chick (*Gallus gallus*), Japanese quail (*Coturnix coturnix japonica*) and white Pekin duck (*Anas platyrhynchos*) were purchased from AA Lab Eggs (Westminster, CA) and incubated at 37°C in a humidified chamber. Eggs were kept in the incubator until they reached embryonic stages appropriate for analysis. For all procedures, we adhered to accepted practices for the humane treatment of avian embryos as described in S3.4.4 of the AVMA Guidelines for the Euthanasia of Animals: 2020 Edition (Leary et al., 2020). Embryos were staged using the Hamburger and Hamilton (HH) staging system, a well-established standard that was originally developed for chicks (Hamburger, 1951). While separate staging systems exist for duck and quail, these embryos can also be staged via the HH system (Ainsworth et al., 2010). Eggs were windowed and external morphological characteristics were used to determine staging. Accurate staging was necessary to ensure proper and consistent timing of collection at HH33 or HH39 for RT-qPCR analysis.

### *Developing* HINTW *PCR primers for quail*

Quail-specific *HINTW* PCR primers were developed using Primer-BLAST (Jian Ye et al., 2012). The input PCR template *Coturnix coturnix japonica Wpkci* (*HINTW*) was retrieved from ENA (AB033881.1) (Hori et al., 2000). The forward primer sequence was 5’-3’ CATTTGAAGATTGTCGGCG (start: 247, stop: 265) and the reverse primer sequence was 5’-3’ CCATGCTCCATCTTTATTGTGC (start: 567, stop: 546). The predicted product length was 321bp, and *in silico* analysis using Primer-BLAST predicted no off-target products.

### Identification of embryo sex using PCR

Genomic DNA was extracted from chick, duck, and quail skin tissue (n=16-20/species) using Extract DNA Prep for PCR (QuantaBio 95091) following the manufacturer’s protocol, and extracts were stored at −20°C. DNA concentration and quality was assessed by a Nanodrop 2000 spectrophotometer (ThermoFisher Scientific, Waltham, MA). All samples were diluted to 200-500ng/μL in ddH2O, and DNA was considered pure at a 260/280 ratio of 1.8±0.1. Reactions were performed in 25μL volumes in a 0.2mL tube containing 12.5μL REDTaq^®^ ReadyMix^™^ PCR Reaction Mix (Sigma-Aldrich R2523), 10.5μL ddH_2_O, 0.5μL forward primer, and 0.5μL reverse primer. All PCR primers are listed in Table 1. Two reactions were run per individual: one reaction targeting the W chromosome (quail *HINTW* primers, chick *HINTW* primers, or *Z43B*-W primers for duck) and one reaction targeting the Z chromosome as an internal and technical control (*Z43B*-Z primers for all species).

Both the quail and chick *HINTW* PCR reactions used the following thermocycling parameters: 98°C for 2 minutes; followed by 35 cycles of 98°C for 10 seconds, 56°C for 30 seconds, and 68°C for 40 seconds; then 68°C for 5 minutes. The *Z43B*-Z and *Z43B*-W PCR reactions used the following thermocycling parameters: 94°C for 15 minutes; followed by 45 cycles of 94°C for 30 seconds, 54°C for 30 seconds, and 72°C for 1 minute; then 72°C for 10 minutes. Products were then loaded in duplicate in 1.2% agarose gels containing MIDORI Green Advance Safe DNA Stain (Bulldog Bio MG04), and the gels were run in 1x TBE running buffer at 100V for 90 minutes. The gel was then imaged using the Azure c400 Gel Imaging System (Azure Biosystems, Dublin, CA). Primer specificity was assessed by comparing actual band size to predicted band size, which are listed in Table 1. Presence of a band in the lane corresponding to a W chromosome primer (quail *HINTW*, chick *HINTW*, or *Z43B*-W for duck) was considered indicative of a female, and presence of a Z chromosome band for the same individual was used as a positive control.

### HINTW *RT-qPCR primer design*

A *HINTW* RT-qPCR primer set effective in chick, duck, and quail was developed using Primer-BLAST (Jian Ye et al., 2012). The input mRNA templates were: chick GenBank NM_204688.2 (Smith et al., 2009), quail ENA AB033881.1 (Hori et al., 2000), and duck ENA AB033883.1 (Hori et al., 2000). The forward primer sequence was 5’-3’ ATGGATTCCGGATGGC and the reverse primer sequence was 5’-3’ CCATGCTCCATCTTTATT. The predicted product length was 272bp for chick and quail and 187bp for duck.

Primer-BLAST was also used to create four of the jaw bone development-related genes analyzed in Figures 3 and 4: *RANK*, *MEPE*, *FGF23*, and *OPN.* The RT-qPCR primers for *CTSK* originate from Houchen et al. 2023, the primers for *MMP13* and *COL1A1* originate from Ealba et al. 2015, and the *RUNX2* primers originate from Ealba et al. 2013. Primer sequences, predicted amplicon sizes, species, and primer sources are listed in Table 1.

### HINTW *and jaw bone development-related gene expression analysis*

Reverse transcription quantitative polymerase chain reaction (RT-qPCR) was used to analyze stage- and species-specific jaw bone development-related mRNA expression, as well as presence or absence of *HINTW* mRNA expression (Bustin et al., 2009). Total RNA was isolated from n=7/stage chick, quail, and duck whole lower jaws at HH33 and HH39 using an RNAeasy column purification kit (Qiagen, Valencia, CA) (Ealba & Schneider, 2013). A Nanodrop 2000 spectrophotometer (ThermoFisher Scientific, Waltham, MA) was used to assess concentration and purity of RNA. RNA was stored at −80°C. Approximately 250ng of total RNA was converted to cDNA in a 20 μL reverse transcription reaction using the Applied Biosystems™ High-Capacity cDNA Reverse Transcription Kit (ThermoFisher Scientific, Waltham, MA). The reaction involved: 25°C for 10 minutes; 37°C for 120 minutes; 85°C for 5 minutes; then 4°C hold in a C1000 Touch Thermal Cycler (Bio-Rad, Hercules, CA). cDNA was stored at −20°C.

RT-qPCR was performed with a StepOnePlus^™^ Real-Time PCR System (ThermoFisher Scientific, Waltham, MA). Forward and reverse primers, 2 μL of cDNA, RNase-free dH2O, and iQ SYBR-Green Supermix (Bio-Rad, Hercules, CA) were manually mixed in a 20 μL reaction to amplify the cDNA of interest. Samples were run in triplicate on white hard-shell 96-well plates (Bio-Rad, Hercules, CA). The protocol was: 95°C for 3 minutes; then 40 cycles of 95°C for 10 seconds, 60°C for 30 seconds, and a plate read; followed by 95°C for 10 seconds; finished by melt curve for 60-90°C for 5 seconds at each 0.5°C with a plate read. Melt curves were analyzed to confirm specificity of primer. RT-qPCR products that were amplified after 35 cycles were considered false positives. Following RT-qPCR, products were run on a 1% agarose gels containing MIDORI Green Advance Safe DNA Stain (Bulldog Bio MG04), and the gels were run in 1x TBE running buffer at 100V for 90 minutes. The gel was then imaged using the Azure c400 Gel Imaging System (Azure Biosystems, Dublin, CA). Predicted amplicon size was compared to band size; this was used as a primer specificity check in addition to melt curve analysis.

All RT-qPCR primer sequences and predicted amplicon sizes are listed in Table 1. Primers were designed for use in both quail and duck using Primer-BLAST software (J. Ye et al., 2012). Expression levels of all jaw development-related genes were normalized to expression of the reference gene glyceraldehyde 3-phosphate dehydrogenase (*GAPDH*). Expression was checked to make sure amplification efficiencies were equal among samples. Fold changes were calculated using the delta-delta C(t) method (Livak & Schmittgen, 2001). To assess relative fold changes between stages, expression at HH39 is relative to expression at HH33.

### Statistical Analysis

RT-qPCR data are shown by mean ± standard deviation with all data points shown. Male HH39 gene expression versus female HH39 gene expression was compared by an unpaired two-tailed Student’s *t* test with a significance cutoff of p<0.05.

## Acknowledgments

We thank T. Dam at AA Lab Eggs for chick, quail, and duck eggs; E. Thompson for assisting with background research and literature review; E. Lee and M. Rawoot for contributing to methods development; and J.M. Scott for statistical analysis recommendations and feedback.

## Author Contributions

E.E.B. conceived and coordinated the project. C.J.H and M.B.G. performed the experiments and compiled the data. C.J.H, M.B.G., and E.E.B. designed the experiments, established the methods, analyzed the data, co-wrote, and critically revised the manuscript. All authors have read and approved of the manuscript and agree to be accountable for all aspects of the work.

## Data Availability Statement

The data that support the findings of this study are available from the corresponding author upon reasonable request.

